# Effects of childhood hearing loss on the subcortical and cortical representation of speech

**DOI:** 10.1101/2024.02.22.581639

**Authors:** Axelle Calcus, Stuart Rosen

## Abstract

Little is known about the effects of childhood mild-to-moderate sensorineural hearing loss (MM HL) on the function of the auditory pathway. We aimed to examine the effect of childhood MM HL and the benefit of frequency-specific amplification on both subcortical and cortical auditory processing, and to relate it to speech-perceptual abilities. We recorded subcortical and cortical responses to speech syllables in nineteen children with congenital MM HL (unamplified and amplified), and sixteen children with typical hearing (unamplified sounds only). Speech perception was measured behaviourally. Congenital HL led to smaller subcortical and cortical responses to unamplified speech sounds. There was a significant benefit of amplification on subcortical and early, but not late, cortical responses, with some effects differing across age. No relationship was found between the neural and behavioural measures. Childhood MM HL affects both subcortical and cortical processing of speech. Amplification mostly benefits subcortical processing of speech in younger children. Childhood HL leads to functional changes in the processing of sounds, with amplification differentially affecting subcortical and cortical levels of the auditory pathway.

## Introduction

Animal studies have shown biological and physiological alterations in the properties of neurons in the cochlea, subcortex, and cortex following even a temporary, mild-to-moderate (MM) hearing loss (HL; Caras & Sanes, 2015; Mowery et al., 2015). Similar changes have been documented in humans with congenital deafness (Daly et al. 1976; Ponton et al. 2000; Sharma et al. 1997), or in older adults with age-related MM HL (for a review see Peelle & Wingfield 2016). Nowadays, most individuals with MM HL who seek treatment receive auditory stimuli via hearing aids, which amplify the incoming auditory signal. To date, the bulk of studies investigating the effects of MM HL on the functional integrity of the auditory pathway (and the potential benefit of hearing aid amplification) has focused on older adults. However, congenital hearing loss affects up to 2 live births in 1000 (Lieu et al. 2020), milder degrees of loss being more prevalent yet more often overlooked. Here, we evaluate the effect of childhood MM HL and the benefit of amplification on simultaneously recorded subcortical and cortical responses to speech sounds.

In the auditory system, speech sounds travel from the cochleae to the auditory cortices, undergoing increasingly complex levels of processing. At the level of the cochleae, sounds undergo a frequency analysis. The output of each channel can be separated into their relatively slowly fluctuating modulated signals known as the envelope (ENV), imposed upon a rapidly varying carrier waveform, known as the temporal fine structure (TFS). Both ENV and TFS cues are thought to be important in decoding speech. ENV cues support the robust identification of speech in quiet (Shannon et al. 1995), whereas TFS cues are thought to be especially useful in noisy backgrounds (Qin & Oxenham 2003; Zeng et al. 2005). Indeed, individuals with typical hearing (TH) typically benefit from the TFS when they glimpse into the “dips” (i.e., local improvements in SNR) of a fluctuating background noise (Hopkins & Moore 2009; Cooke 2006). Therefore, speech intelligibility is much better in the presence of fluctuating than steady background noise, a phenomenon termed “masking release” (Miller & Licklider 1950; Festen & Plomp 1990). However, adults with HL show reduced ability to use TFS (Lorenzi et al. 2006), a difficulty that limits the amount of masking release they experience (Hopkins et al. 2008; Strelcyk & Dau 2009). Similarly to adults, children and adolescents with MM HL experience poorer speech intelligibility in noise (Crandell 1993; Lewis et al. 2015; Goldsworthy & Markle 2019). They show reduced sensitivity to TFS compared to children/adolescents with TH (Halliday et al. 2019), which likely limits masking release of speech in the presence of a fluctuating background noise, even when they wear hearing aids (Brennan et al. 2016).

At the subcortical level of the auditory pathway, the frequency following response (FFR) reflects the periodicity of speech sounds (Kraus et al. 2017). Adding the neural responses to speech sounds presented in alternating polarities is thought to enhance the ENV of the response (henceforth FFR_ENV_, here used as an index of fundamental frequency encoding), whereas subtracting them is considered to enhance the TFS (henceforth FFR_TFS_, an index of spectral harmonic encoding (Aiken & Picton 2008)). Although the primary generator of the FFR appears to vary according to the fundamental frequency (F0) of the input stimulus, the current view is that the FFR emerges from multiple generators, including the cochlear nucleus, inferior colliculus and thalamus (Bidelman 2018; Coffey et al. 2019). Early studies suggested that the auditory brainstem reached maturity within the first 18 months of life (Hecox & Galambos 1974). However, more recent studies indicate that the FFR evoked by speech sounds keeps maturing well into the childhood years (Krizman et al. 2015; Krizman et al. 2019).

At the cortical level, components of the late auditory event-related potentials (LAERs) arise which are thought to reflect the initial detection (P1, N1, and P2), classification and discrimination (P2 and N2) of auditory stimuli, reflecting processing for auditory events ranging from isolated sounds to complex auditory scenes (Alain & Tremblay 2007). LAERs are thought to be generated in the auditory cortex (for a review, see Eggermont & Ponton, 2002), with additional contributors from non-auditory (e.g., prefrontal, premotor) areas (Scherg et al. 1989). The mismatch negativity (MMN) is thought to reflect the discrimination of a deviant sound, and is typically evoked by a comparatively rare deviant in a stream of repeated standards (i.e. an oddball paradigm; for a review see Näätänen et al. 2012). The MMN is thought to have several additional generators, including bilateral temporal cortex, right inferior frontal gyrus, and bilateral frontal and centro-parietal regions (e.g., Alho 1995; Rinne et al. 2000; Zhang et al. 2018). In addition to distinct topographies, both LAERs and MMNs show distinct developmental trajectories. In typical development, P1 and N2 responses are thought to decrease in amplitude as children grow older, while N1 and P2 progressively appear in childhood, growing in amplitude until adolescence (Bishop et al. 2007; Mahajan & McArthur 2012). MMN amplitude is thought to increase throughout childhood (Wetzel et al. 2006), but to be stable from around 10 years of age to adulthood (Mahajan & McArthur 2015; Mueller et al. 2008) - although some studies report a prolonged increase in MMN amplitude throughout adolescence (Bishop et al., 2011; Oades et al., 1997).

Both subcortical and cortical levels of the auditory pathway thus appear to follow a protracted trajectory, maturing until late adolescence. How MM HL affects the development of subcortical and cortical speech processing remains largely unknown. So far, studies have focused on *either* subcortical *or* cortical levels of processing, only providing fragmented information about the physiology of the auditory pathway in children/adolescents with MM HL. Yet using the right set of recording/analysis parameters, it is now possible to obtain subcortical and cortical responses during the same recording session (Krishnan et al. 2012; Bidelman, 2015). Therefore, the first aim of this project was to evaluate the effect of childhood hearing loss on simultaneously recorded subcortical and cortical levels of processing. Noteworthily, the existing literature on this topic can be classified according to the strategy used to manage intensity levels across listener groups.

### Neural processing of speech presented at fixed intensity levels across groups with and without HL

When studying the effect of hearing loss on auditory processing, researchers often choose to investigate the unaided performance of individuals with HL. Stimuli are thus presented at a fixed intensity level for all individuals, with HL or not. Adult studies using this strategy to evaluate the effect of HL on subcortical processing have led to contradictory findings (for a review, see Jacxsens et al. 2024). While some authors reported smaller FFR_ENV_ and FFR_TFS_ in adults with HL than adults with TH (Plyler & Ananthanarayan 2001; Ananthakrishnan et al. 2016), others did not show such group differences (Anderson et al. 2013; Presacco et al. 2019). Note that a possible explanation for this discrepancy lies in the age-matching across groups (Anderson et al. 2013; Presacco et al. 2019), or lack thereof (Plyler & Ananthanarayan 2001; Ananthakrishnan et al. 2016). To our knowledge, no study has focused on the effects of childhood MM HL on subcortical processing of speech, using fixed intensity stimuli across groups of children/adolescents.

At the cortical level, studies using fixed intensity levels across groups suggest that childhood MM HL leads to delayed and/or smaller LAERs evoked by speech or pure tones (Calcus et al. 2019; Ji et al. 2023). Developmental changes following MM HL were also apparent in the MMN: While 8-11 year old children with MM HL showed age-appropriate MMN responses to sounds, this was no longer the case in 12-16 year old adolescents with MM HL, as shown both cross-sectionally and longitudinally (Calcus et al. 2019). This suggests that some effects of childhood MM HL might only reveal themselves in adolescence, highlighting the need to take age into account in developmental studies.

### Neural processing of speech presented at similar sensation levels across groups

A common limitation of the above-mentioned studies using fixed intensity levels for all participants is that individuals with HL would have perceived the stimuli at reduced sensation levels (SL) relative to controls. Therefore, some researchers have attempted to equate for the effects of peripheral hearing loss by presenting listeners with TH and those with SNHL are presented with sounds at different intensities – more intense for individuals with SNHL (either by simulating the frequency-specific gain provided by individuals’ hearing aids, or by recording the neurophysiological response while listeners wear hearing aids). With such adjustments for HL, adults with acquired MM HL show larger FFR_ENV_ to an amplified speech stimulus compared to an unamplified one (BinKhamis et al. 2019; Easwar et al. 2015) and in one study, than the FFR_ENV_ evoked by unamplified speech presented to TH listeners (Anderson et al. 2013). A similar benefit of amplification on the FFR_ENV_ evoked by the speech token /su∫i/ was recently reported in 18 children with HL aged 6 to 17 years, compared to the unamplified stimulus (Easwar et al. 2023). Note that older adults with acquired MM HL did not show a significant benefit of amplification on FFR_TFS_ (Anderson et al., 2013). To our knowledge, the effect of amplification and childhood HL on FFR_TFS_ has remained unexplored so far.

At the cortical level, studies looking at the effect of childhood HL on LAERs by accounting for the effects of peripheral HL have led to contradictory findings. Pilot data from a small sample of 5 children aged 2 to 6 years show age-appropriate P1 and N2 evoked by speech stimuli when the children wore their hearing aids (Martinez et al. 2013). Another study focused on older, unaided children (9-12 years) with MM HL, to whom speech stimuli were presented on average 18 dB greater than in children with TH (Koravand et al. 2013). Note that here, the amplification provided to the group with HL was not frequency-specific – unlike hearing aids. Results indicate age-appropriate P1 and MMN responses in children with MM HL, but smaller (but not later) N2 responses compared to children with TH. However, 5 to 8 year-old children who wore their hearing aids (n = 7) showed significantly smaller LAERs evoked by complex tones than age-matched children with TH (Engström et al. 2021). Interestingly, when re-tested three years later, the same children with HL showed age-appropriate LAERs. At both time-points, MMN amplitudes were age-appropriate in children with MM HL. To our knowledge, no study has directly investigated the intra-individual benefit of amplification on the cortical processing of speech in children/adolescents with MM HL.

### Relationship between speech processing and perception

A main stake of auditory neurophysiological studies is to relate neural processing to speech perception in a variety of contexts and clinical populations. Because the FFR reflects encoding of F0 and lower harmonics of the signal, that are particularly helpful in adverse listening environments, it is thought to relate to speech perception in noise. Better speech perception in noise has been associated with subcortical processing of complex sounds in quiet, as observed in children (Anderson et al. 2010; Hornickel et al. 2009), young (Bidelman & Momtaz 2021) and older adults with TH (Mai et al. 2018; Mai & Howell 2023); but see Presacco et al., 2016 for a study that did not find such an association). In fact, FFR_ENV_ evoked by amplitude-modulated tones presented in quiet even seems to predict sentence understanding in noise in adults with TH (Mepani et al. 2021). Recent evidence suggests that FFR_ENV_ (evoked by speech in quiet) correlates with speech perception in noise in unaided (but not in aided) 4 to 9 year-olds with congenital mild to moderate HL (van Hirtum et al. 2023). Whether subcortical processing relates to speech perception in noise in older children/adolescents with congenital HL remains an open question.

Cortical processing of sounds is also thought to relate to behavioural speech perception, mostly in quiet. In 4-month old children with TH or HL, components of the LAER relate to parents’ reports of auditory functional performance (Ching et al. 2023). In adults, the MMN is thought to relate to perceptual speech discrimination, at least at the group level (for a review, see Näätänen 2001). However, no clear relationship has emerged between MMN amplitude and speech discrimination in children with TH so far (Sharma et al. 2006; Jost et al. 2015; Shafer et al. 2005). To our knowledge, it remains unclear whether cortical EEG measures relate to speech perception in children/adolescents with congenital HL.

### Study objectives

The main goal of this study was twofold: to evaluate (i) the effect of childhood HL and (ii) the benefit of amplification on both subcortical and cortical auditory processing. (i) So far, no study has investigated the effect of childhood MM HL on subcortical processing presented at fixed intensity levels across age-matched groups. Additionally, results from (older) adult studies have led to contradictory results. Therefore, this study will be a first investigation of the effect of MM HL on the processing of unamplified sounds throughout development. At the cortical level, we predict smaller/later LAER and MMN responses in children/adolescents with HL than those with TH, with a possible interaction between age and group on the MMN. (ii) Regarding the second objective, we expect larger FFR_ENV_ in children with HL when sounds are amplified than unamplified. At the cortical level, results from the literature are mixed, but have never compared LAER/MMN with and without amplification in the same children with HL. We will thus study the effect of amplification on the cortical responses evoked in children/adolescents with HL.

Given that subcortical and cortical processing have both been related to speech perception, an exploratory objective was (iii) to investigate how these neurophysiological responses relate to speech intelligibility in quiet and in noise.

## Methods

### Participants

Thirty-five children aged 8 to 16 years were included in the study. Nineteen (8 female) had a diagnosis of mild-to-moderate SNHL (HL group) and sixteen (8 female) were age-matched children with typical hearing (TH group). All participants were monolingual British English natives and obtained a nonverbal IQ (NVIQ) score of at least 85 (Block Design subtest of the Weschler Abbreviated Scale of Intelligence). No child had any known medical, psychological, neurological or developmental disorders other than SNHL. Individual air-conduction audiometric thresholds were measured bilaterally in all children at octave frequencies from 0.125 to 8 kHz using an Interacoustics AC33 audiometer. The study was conducted with the verbal assent of the participants, the written consent of their parents, and was approved by the UCL Research Ethics Committee (2109/004). It was performed in accordance with the Declaration of Helsinki.

**Figure 1:**
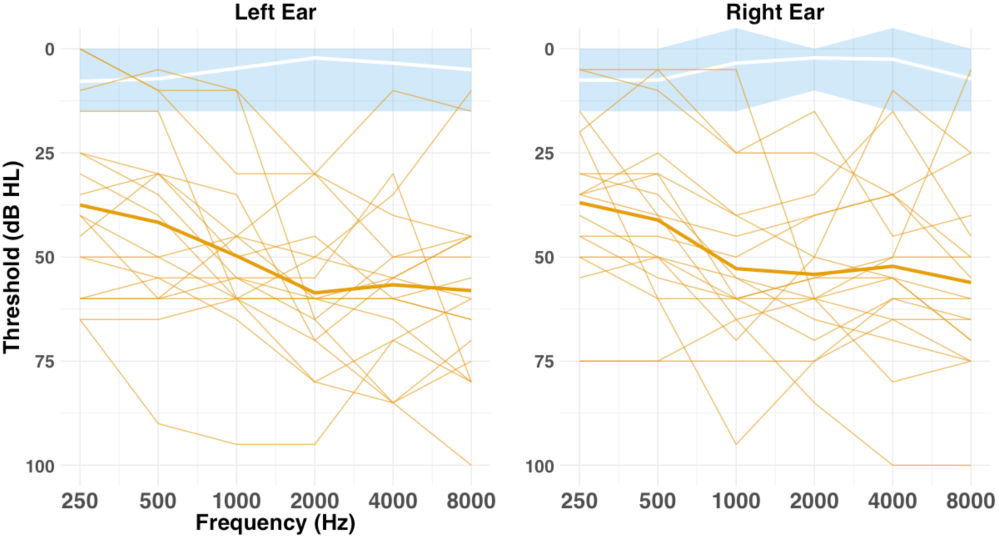
Pure-tone air-conduction audiometric thresholds for children with HL (orange) and children with TH (blue and white). Audiometric thresholds are shown across octave frequencies from 0.25 to 8 kHz in the left and right ears. Individual thresholds for the HL group are shown as lines, and the group mean is shown as a bold line. Mean thresholds for the TH group are marked in white, with the shaded blue area representing the range for the TH group.

Participants in the HL group had better-ear pure tone average (BEPTA) thresholds of 21-70 dB HL across 0.25-8 kHz (see Figure 1), with on average 7 dB difference in PTA between the ears. All children in the HL group were spoken language users (i.e., non-signers), and sixteen had prescription hearing aids, but these were not used in this study. Children in the HL group were individually matched in age (± 6 months) to those in the TH group, and mean age did not differ significantly between the two groups (see Table 1). None of the children in the TH group had a known history of hearing loss (including otitis media with effusion), educational difficulties, or speech and language problems (based on parent/guardian report). All had mean PTA thresholds ≤ 20 dB HL across octave frequencies 0.25-8 kHz in both ears and obtained thresholds no higher than 20 dB HL at any given frequency.

Group comparisons are presented in Table 1. Note that some children did not complete all behavioural and electrophysiological tasks. With respect to the behavioural tasks, two children (1 TH and 1 HL) did not perform the speech identification in noise tasks. Due to technical difficulties, two children (1 TH and 1 HL) did not perform the speech identification in a fluctuating noise task. Two children with HL did not perform the speech discrimination in quiet task. With respect to the EEG measurement, one child (HL) was only tested in the amplified, but not the unamplified EEG condition. The sampling frequency of the EEG recording was accidentally set to 2048 Hz for another child (HL). For this child, we could obtain the cortical measures, but not the subcortical measures. In all these cases, results were treated as missing data.

**Table 1:** Mean (SD) participant characteristics for the HL and TH groups. Comparisons are independent-samples *t*-tests. Effect sizes are Cohen’s *d*. Significant effects are shown in boldface.

| Variable | TH<br>(n = 15) | HL<br>(n = 19) | *t*(df) | *p*-val | Effect<br>size | 95 % CI |
| --- | --- | --- | --- | --- | --- | --- |
| Age (years) | 12.6 (2.43) | 12.4 (2.44) | 0.26<br>(32.1) | .795 | 0.08 | [-1.46, 1.89] |
| **Nonverbal IQ** | 119.1 (12.6) | 102.5 (12.5) | 3.83<br>(31.44) | **<.001** | 1.32 | [-25.4, -7.7] |
| **Maternal education** | 20.9 (2.59) | 18.3 (2.78) | 2.76<br>(30.99) | **.009** | 0.96 | [-4.49, -0.67] |
| **BEPTA threshold**<br>**(dB HL)** | 3.7 (2.87) | 45.0 (13.30) | -13.18<br>(19.97) | **<.001** | 4.12 | [-47.86, -34.78] |
| Age of HL detection<br>(years) | - | 3.6 (3.03) |  |  |  |  |

### Data acquisition and pre-processing

Stimuli were digitised /bɑ/ (standard) and /dɑ/ (deviant) syllables (see Figure 2) originally spoken by a native British female speaker, and identical to those used in Halliday et al., 2019; Calcus et al., 2019; Bishop et al., 2011. The intonation contours were made monotone at 220 Hz using Praat (Boersma & Weenink, 2001) and the final stimuli were RMS equalised with GoldWave (Craig, 2008). Vowel formant frequencies were approximately 800, 1340 and 2700 Hz for formants F_1_, F_2_ and F_3_ respectively, although there were small differences between the two syllables as the stimuli were based on natural utterances. Both syllables had rising F_1_s over ∼50 ms with a much larger transition for the /dɑ/ (∼230 Hz) than the /bɑ/ (∼115 Hz). The phonetic contrast was cued primarily by differences in the properties of the initial release burst (more intense for the /dɑ/, especially at higher frequencies) and on the formant transitions into the vowel. No F_2_ transition was present for the /bɑ/ but the /dɑ/ had a significant falling transition for F_2_ of ∼350 Hz.

Each condition consisted of a total of 1320, 175-ms stimuli, presented in a passive oddball paradigm (deviant probability: 10%). Half of all stimuli were presented in positive polarity and the other half in negative polarity. The order of presentation of each polarity was randomized. The inter-stimulus interval was jittered randomly between 275 and 375 ms to limit neural adaptation (Malmierca et al., 2014).

**Figure 2:**
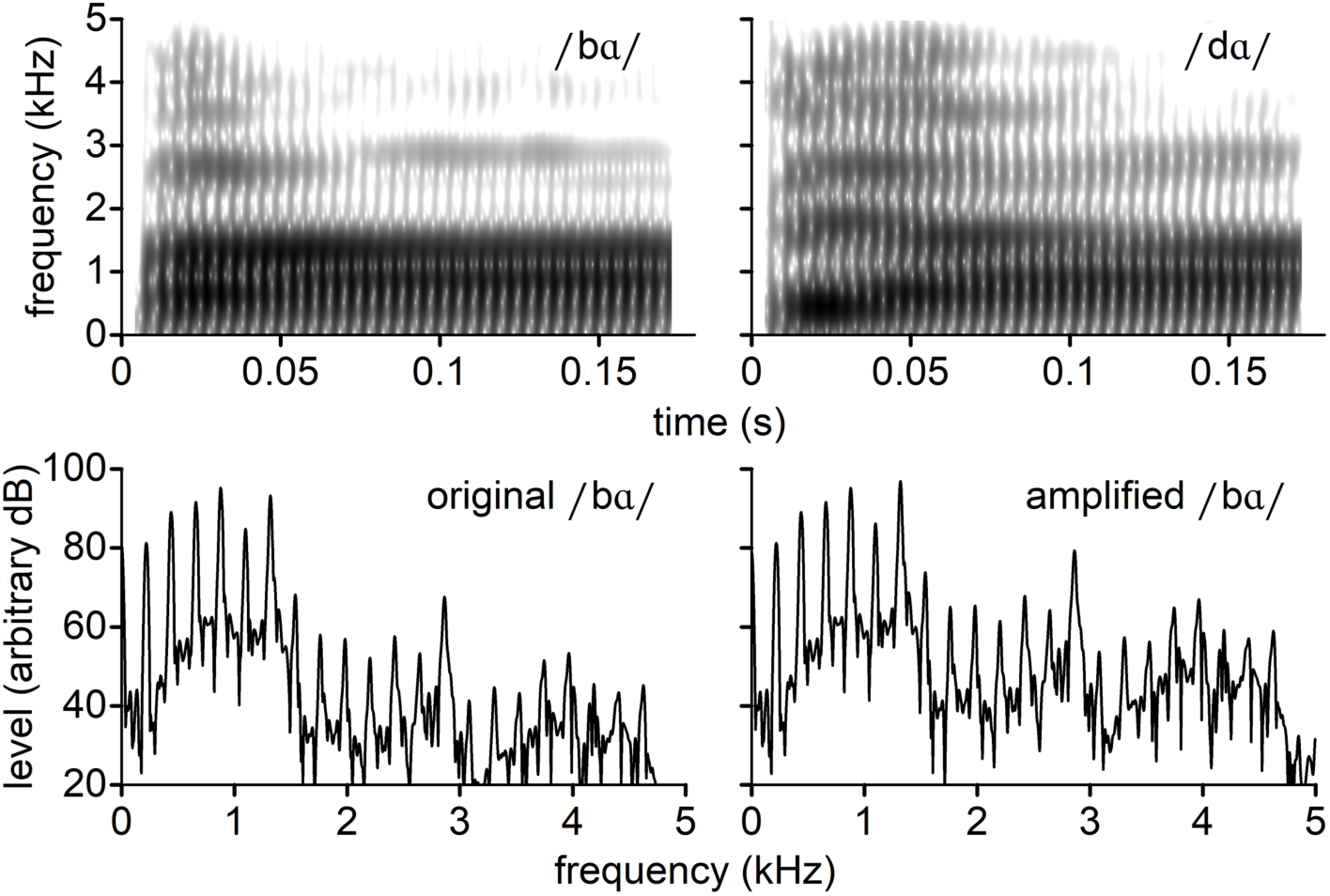
Top row: Spectrograms of the standard and deviant speech syllables in their unamplified form. Bottom row: Spectra of the vocalic portion (20 to 170 ms) of the standard syllable /bɑ/ in both unamplified and NAL-R amplified versions. Children with TH were only presented with unamplified versions of the stimuli. Children with HL were presented with both versions of the stimuli, with the amplification being dependent upon each individual’s audiogram. For visualisation purposes, the amplified stimulus here was generated based on the average audiogram of the HL group. It can clearly be seen that, as expected, the level of the harmonics below ∼2 kHz is unchanged because there is minimal hearing loss at those frequencies, with amplification of the higher harmonics in the frequency region in which there is substantial HL.

Children with TH were presented with only one condition, consisting of unamplified stimuli that were presented binaurally at 70 dB SPL through shielded insert earphones (ER-2, Etymotic Research) – henceforth referred to as children with TH_U_. Children with HL undertook two conditions - unamplified and amplified – presented in counter-balanced order (henceforth: HL_U_ and HL_A_). Amplification used the National Acoustic Laboratory-Revised algorithm (NAL-R, Byrne & Dillon, 1986) to provide a frequency-specific gain (without compression), based on each individual child’s audiogram. This strategy minimizes the risk of potential confounds related to sound-field testing as well as potential inter-individual variability in the hearing aid processors of individual children (Billings, 2013). Stimuli in the amplified condition were limited in intensity such that they did not exceed 85 dB SPL. Each condition lasted approximately 20 minutes, with three short breaks offered every 5 minutes. During the recording, participants were seated comfortably in a shielded, sound-attenuated booth, and watched a silent movie.

EEG was recorded using a BioSemi ActiveTwo system at a sampling rate of 8192 Hz, from 32 scalp electrodes in the standard 10/20 configuration (Jasper, 1958). Additional electrodes were placed on each mastoid and recordings were re-referenced offline to the average of activity at the mastoid electrodes. Electrode offsets were consistently < 50 mV.

### Subcortical response analysis

Subcortical response analyses were performed only on the standard (/bɑ/), to avoid the confound of small spectral differences between the standard and the deviant syllables. 1188 epochs (standard probability: 90%) were obtained by applying a band-pass filter from 90 to 1600 Hz (zero-phase, finite impulse response, -6dB/octave) to the EEG data evoked by the standard syllables (/bɑ/) at Cz, and extracting data 25-175 ms relative to trigger onset, which covers the duration of the steady portion of the vowel. Trials exceeding ± 100 µV were rejected before averaging. There was no significant difference in the proportion of trials that were rejected across groups [TH_U_: 18.5%; HL_U_: 19.1%; HL_A_: 20.3%; *F*(2, 47) = 0.723, *p* = 0.491, *η^2^* = 0.03]. The averaged responses to positive and negative polarities were then added together to enhance the envelope of the response (FFR_ENV_), and subtracted from each other (i.e. positive minus negative polarity) to enhance the TFS of the response (FFR_TFS_). Spectral amplitudes were calculated using a fast Fourier transform (FFT), after applying zero-padding to the sampling rate to obtain an FFT with a resolution of 1 Hz. Spectral peaks at F0 (220 Hz) and harmonics (H2: 440 Hz; H3: 660 Hz; H4: 880 Hz; H5: 1120 Hz; and H6: 1320 Hz) were computed as the maximum amplitude within an 8-Hz bin centred on F0 and subsequent harmonics, following visual inspection of the grand average subcortical response, and in line with previous studies (e.g. Schoof & Rosen, 2016). Neural background noise amplitudes were calculated by taking the mean amplitude of the fast Fourier transform across 100-Hz windows surrounding each spectral peak (50 Hz on each side, excluding the ±70 Hz adjacent bins).

### Cortical response analysis

Evoked potentials considered to be cortical in origin were obtained by band-pass filtering the EEG recorded at electrode Fz between 0.5 Hz and 35 Hz (zero-phase, finite impulse response, -6dB/octave) and creating epochs from -100 ms to 500 ms relative to each stimulus onset time. 1320 epochs (1188 standards and 132 deviants) were baseline corrected using the mean value from -100 to 0 ms. Trials exceeding ± 100 µV were rejected before averaging. There was no significant difference in the proportion of trials that were rejected across groups [TH_U_: 12.7%; HL_U_: 13.4%; HL_A_: 15.75%; *F*(2, 50) = 0.43, *p* = 0.654, *η^2^* = 0.01].

P1 and N2 grand average latency were quantified by means of the half-area latency measure of the standard waveform recorded in the TH group (Zhang & Luck, 2023), a computation that is less sensitive to high-frequency noise than traditional latency peak measures (Finke et al., 2016; Luck, 2014). The window for computation of the area under the curve of the P1 ranged from 55 to 135 ms and from 250 to 350 ms for the N2, based on visual inspection of the TH grand average. P1 and N2 amplitudes were then computed for each participant and condition as the mean amplitude within a 40-ms window centred around the half-area latency (P1: 96 ms post-stimulus onset; N2: 329 ms post-stimulus onset). P1 and N2 latencies were computed for each participant and condition as the half-area latency for the local peak to appear, within the 40-ms window centred around the TH grand average evoked by standards. Note that N1 and P2 were not analysed because they could not be reliably identified in the younger participants, which is likely due to a protracted maturation of these responses until adolescence (Bishop et al., 2011; Sussman et al., 2008).

The MMN was computed as the differential wave obtained by subtracting the neural response evoked by the standard to that evoked by the deviant, for each participant and condition. The MMN was then statistically assessed in two ways. First, to determine whether there was evidence for neural discrimination of speech deviants, the presence of an MMN was evaluated for the three combinations of group and amplification conditions. Point-to-point comparison of the differential wave amplitudes was performed to determine the latency period over which the waveforms were significantly smaller than zero, if any. One-sided *t*-tests were computed within the 100-500 ms post-stimulus-onset time window which was identified as the region most likely to contain the MMN by visual inspection of the grand average waveforms. Because adjacent points in the waveform are highly correlated, potentially leading to spurious significant values in short intervals, an MMN was considered present when *p* < .01 (one-tailed) for more than 20 ms at adjacent time-points (Kraus et al., 1993; McGee et al., 1997). Half-area MMN latency was computed on the differential waveform recorded in the TH group. MMN amplitude for each participant and condition were then computed as the mean amplitude in a 100-ms window centred around the grand averaged differential waveform half-area latency peak observed in children with TH (at 426 ms post-stimulus onset; Zhang & Luck, 2023). MMN individual latencies were computed as the half-area latency within a 100-ms window centred around the grand average differential waveform of the children with TH.

### Speech discrimination in quiet

The speech discrimination in quiet task was the same as that used in previous studies (Halliday et al., 2017; 2019). The two endpoints of the continuum were based on the same digitised /bɑ/ and /dɑ/ used in the electrophysiological recordings. A continuum of 100 stimuli was then constructed (including the endpoints), using the morphing capabilities of the programme STRAIGHT (Kawahara et al., 1999). The task was delivered via a child-friendly computer game, incorporating an adaptive, three-interval, three-alternative, forced-choice paradigm. Stimuli were presented binaurally via headphones (Sennheiser 25 HD) at a fixed intensity (70 dB SPL), on a tablet, meaning identical levels presented in the two groups. Children were required to detect the odd-one-out, where two of the intervals corresponded to the /bɑ/ end of the continuum, and the third, target stimulus fell somewhere along the rest of the continuum. Participants received visual feedback regarding the accuracy of their responses. A three-down, one-up procedure was used to select, on each trial, the appropriate target sound from a continuum, tracking a performance level of 79.4% correct (Levitt, 1971). This was preceded by an initial one-down, one-up procedure until the first reversal (Baker & Rosen, 2001). The initial step size was 15 stimulus places along the continuum, which reduced to five after the first reversal. The threshold was the arithmetic mean of the last four reversals in direction of the adaptive track, expressed as the stimulus number (0-99) of the target stimulus along the continuum. Therefore, smaller numbers indicate better performance. The task was preceded by five practice trials which contained standard-target difference that had previously been deemed suprathreshold for adults with TH (Halliday et al. 2019).

### Speech perception in noise

The rationale for choosing the stimuli for this task was to provide the most direct comparison to the neurophysiological stimuli, which were speech syllables. Another advantage of syllables is that they minimise the role of linguistic and cognitive factors on auditory performance, allowing to focus more on lower level auditory properties. Speech reception thresholds (SRTs) were thus measured adaptively for 13 consonants presented in vowel-consonant-vowel (VCV) logatomes, e.g. /imi/, /apa/. Vowels were /i/, /ɑ/, and /u/, and consonants were /p/, /k/, /b/, /d/, /g/, /f/, /m/, /n/, /l/, /v/, /w/, /y/, and /z/. Both the vowels and consonants varied from trial to trial, but only the response for the consonant was scored. Stimuli were recorded by a native British English female speaker in an anechoic chamber and were digitized via a 16-bit analogue-to-digital converter at a 44.1-kHz sampling frequency. The logatomes were presented together with two types of background noise: steady (i.e. unmodulated, but see Stone & Moore, 2014) or fluctuating in amplitude sinusoidally at 8 Hz, which corresponds to a syllabic modulation rate (Greenberg et al., 2003). The modulation depth was fixed at 1.0 and the starting phase was randomized between 0° and 360° on each trial. The noise was shaped on-and-off using a raised-cosine with 50-ms rise-fall times.

Participants were seated in a quiet room and the stimuli were presented binaurally over Etymotics ER-2 earphones at an overall level of 70 dB SPL, again meaning identical levels presented in the two groups. They were asked to repeat the logatomes as best as they could. The experimenter scored the participants’ responses using a graphical interface which showed the 13 possible consonants. No feedback was provided.

The expected wide variability in intelligibility across consonants at a given SNR would not allow their reliable use in a simple adaptive procedure. Therefore, SRTs for each combination of consonant and vowel were estimated from a prior study of young adults with TH (unpublished), allowing the calculation of adjustment factors in SNR to give all stimuli the same nominal SRT (Plomp & Mimpen, 1979). These factors were then used during the adaptive procedure to adjust the presented SNR. The signal-to-noise ratio (SNR) was varied adaptively by varying the level of the target and fixing the level of the noise at 70 dB SPL. The first logatome was presented at a nominal SNR of -10 dB (taking into account that token’s adjustment factor). Depending on the participant’s response (correct/incorrect), the SNR of the second logatome was decreased/increased by 10 dB. Again, depending on the participant’s response (correct/incorrect), the SNR of the third logatome was decreased/increased by 6 dB. For each subsequent logatome, the SNR increased/decreased by 2 dB for incorrect/correct responses, respectively. Measurement stopped after either 12 reversals or 78 trials, whichever occurred first. The SRT was computed by taking the mean SNR (dB) across all track reversals.

### Speech perception in quiet

As a baseline and familiarisation for the speech perception in noise task, we also included a speech perception in quiet task. Percentage correct responses were measured for 78 consonants presented at a fixed intensity of 70 dB SPL, using the same VCV logatomes and procedure for recording the participants’ responses as those used in the speech perception in noise task.

### Statistical analyses

Regarding the subcortical data, we first ran an outlier detection algorithm. Data points that fell outside the mean ± 3 *SD* (within each combination of group, amplification, FFR component and harmonic) were considered outliers and excluded from further analyses. In total, 2.7% of data points were excluded from the analyses, most of them from the HL group. After excluding outliers, the subcortical data were normally distributed (Kolmogorov-Smirnov test: all *p*s > .05). Note that for both FFR_ENV_ and FFR_TFS_, there were significant differences in variance between groups as evidenced by Levene’s test for equality of variance (both *p*s < 0.001).

Second, paired two-tailed *t*-tests with Bonferroni-Holm correction were conducted separately for each group (TH_U_, HL_U_, HL_A_), FFR component (FFR_ENV_ and FFR_TFS_) and harmonic (F0 to H6) to determine whether the FFR responses were significantly above the neural noise floor. FFR responses are considered present if at least one of the three groups showed a significant difference between target frequency peak and spectral noise floor in each component, at each harmonic. For FFR_ENV_, peaks were significantly larger than the noise floor only at F0 and H2 (all *p*s < 0.001). For FFR_TFS_, peaks were significantly larger than the noise floor at H3, H4, H5 and H6 (all *p*s < 0.05).

Third, a linear mixed model was used to determine whether amplitude of the spectral noise floor differed across groups (TH_U_, HL_U_, HL_A_), FFR component (FFR_ENV_ and FFR_TFS_) or harmonic (F0 to H6). As expected, the only significant predictor was harmonic (*p* < 0.001). In the absence of a significant group effect on the baseline (*p* > 0.10), subsequent analyses were performed on the target frequency peak amplitude.

Regarding the cortical data, we found no outliers (± 3 *SD* from the mean) in the amplitude of either P1 or MMN. The cortical data were normally distributed (Kolmogorov-Smirnov test: all *p*s > .05). For both P1 and MMN amplitudes, there were no significant differences in variances between groups (Levene’s test for equality of variance: both *p*s > .05).

Linear mixed models (lmerTest package of R^92^) were used to evaluate group differences in both cortical (P1, N2 and MMN) and subcortical (FFR_ENV_ and FFR_TFS_) measures. Fixed effect predictors and all their interactions (all orders) were included in the initial model. In all models, the factor listener was used as a random intercept, and random slopes for all the appropriate predictors (harmonic and amplification) that did not lead to convergence issues or inestimable models. Backward stepwise reduction was then used to simplify the models based on F-tests, eliminating insignificant terms, but always keeping lower-order terms that were significant in higher-order interactions. This was done first for the random effects, and then the fixed effects. Only significant results are reported below. Analyses of group differences aimed to answer three main questions.

- First, is there a significant group difference in electrophysiological responses to unamplified speech stimuli? Here, the predictors were group (TH_U_ vs. HL_U_) and age (continuous variable), with harmonic as an additional predictor (FFR_ENV_: F0 and H2; FFR_TFS_: H3 to H6) to model the subcortical data.
- Second, for children with HL only, is there a benefit of amplification on the neural encoding of sounds? Here, the predictors were amplification (HL_U_ vs. HL_A_), BEPTA and age, again with harmonic as an additional predictor for the subcortical data.
- Third, does sound amplification in children with HL lead to electrophysiological responses that are similar to the control group (TH_U_ vs. HL_A_)? Again, the predictors were group (TH_U_ vs. HL_U_) and age, with harmonic as an additional predictor for the subcortical data.

With respect to the speech intelligibility measure in quiet, whose outcome variables was binomial, generalized linear mixed models (logistic regressions) were fitted to the data. Group (TH, HL) and age, as well as their two-way interaction were included in the model as between-subjects factors, with both listener and consonant included as random effects. For speech perception in noise, a linear mixed model was computed with group (TH, HL), noise type (steady, fluctuating), age, and all their interactions as fixed factors, and listener as a random effect. Backward stepwise reduction was then applied, as for the EEG measurements.

Finally, the relationship between subcortical and cortical measures, as well as that between electrophysiological and behavioural measures were investigated using correlations, conducted separately on the TH and HL groups, and separately for unamplified and amplified conditions in the latter group.

**Table 2:**
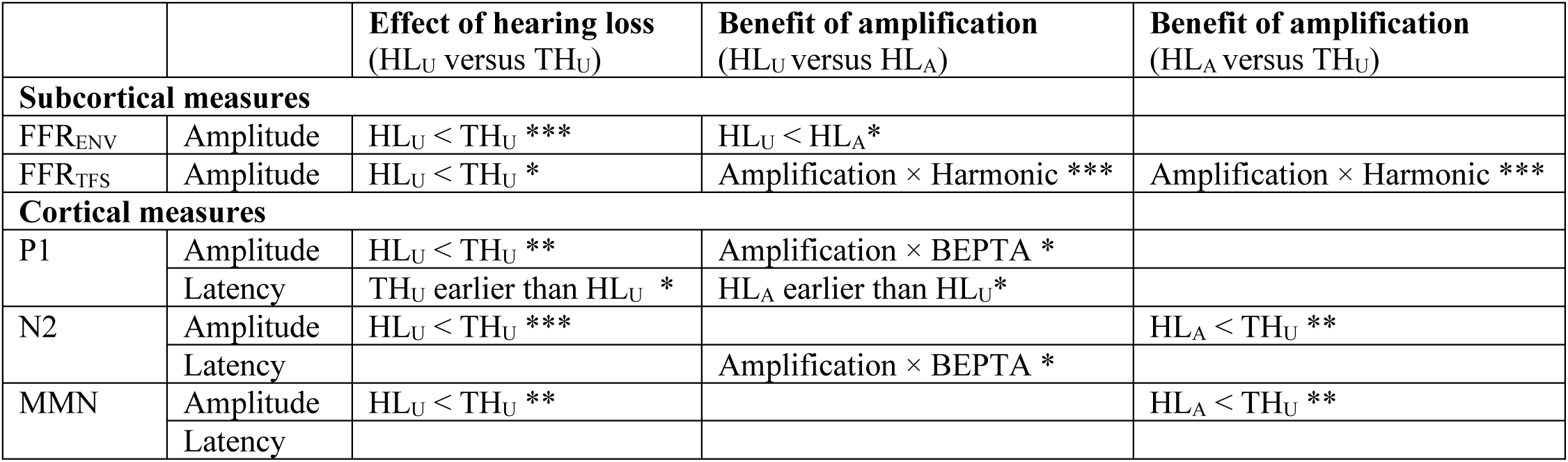
Summary of the main effects of hearing loss (HL_U_ versus TH_U_) and amplification (HL_A_ versus HL_U_) for subcortical, cortical and behavioural measures. * p < 0.05, ** p < 0.01, *** p < 0.001.

## Results

A summary of the main effects of hearing loss (HL_U_ versus TH_U_) and of amplification (HL_A_ versus HL_U_) is provided in Table 2, for subcortical, cortical and behavioural measures.

### 1. Subcortical responses

Figure 3 shows the grand-average time waveforms and spectra of the subcortical responses evoked at electrode Cz by unamplified speech for children with TH (left panels), and unamplified and amplified speech for children with HL (middle and right panels, respectively). Grand-averages were derived from the time waveforms and spectra of the individual waveforms. Scatterplots of the amplitude of the various components are shown as a function of age in Figure 4.

First, group differences in the subcortical processing of unamplified speech were examined (TH_U_ vs. HL_U_; see Supplementary Table 1). FFR_ENV_ were significantly smaller (by 0.082 µV) in children with HL_U_ than in children with TH_U_ (*p* = .001) for both harmonics, with no effect of age. Similarly, FFR_TFS_ were significantly smaller (by 0.028 µV) in children with HL_U_ than in children with TH_U_ (*p* = .015). Note that in neither analysis was there a significant group x harmonic interaction.

Next, the effects of amplification were examined by comparing the subcortical responses of children with HL presented with unamplified versus amplified speech stimuli (HL_U_ vs. HL_A_; see Supplementary Table 2). Overall, FFR_ENV_ were significantly larger (by 0.118 µV) in response to amplified compared to unamplified speech sounds (*p* = 0.011). The amplification × age interaction got excluded during model reduction (*p* = 0.051), yet we observed a marginally significant decreasing benefit of amplification with age on FFR_ENV_ amplitude (*p* = 0.019). With respect to the FFR_TFS_, we observed a significant amplification × harmonic interaction. FFR_TFS_ were significantly larger in response to amplified than unamplified speech sounds at H4, H5 and H6, but not at H3.

**Figure 3:**
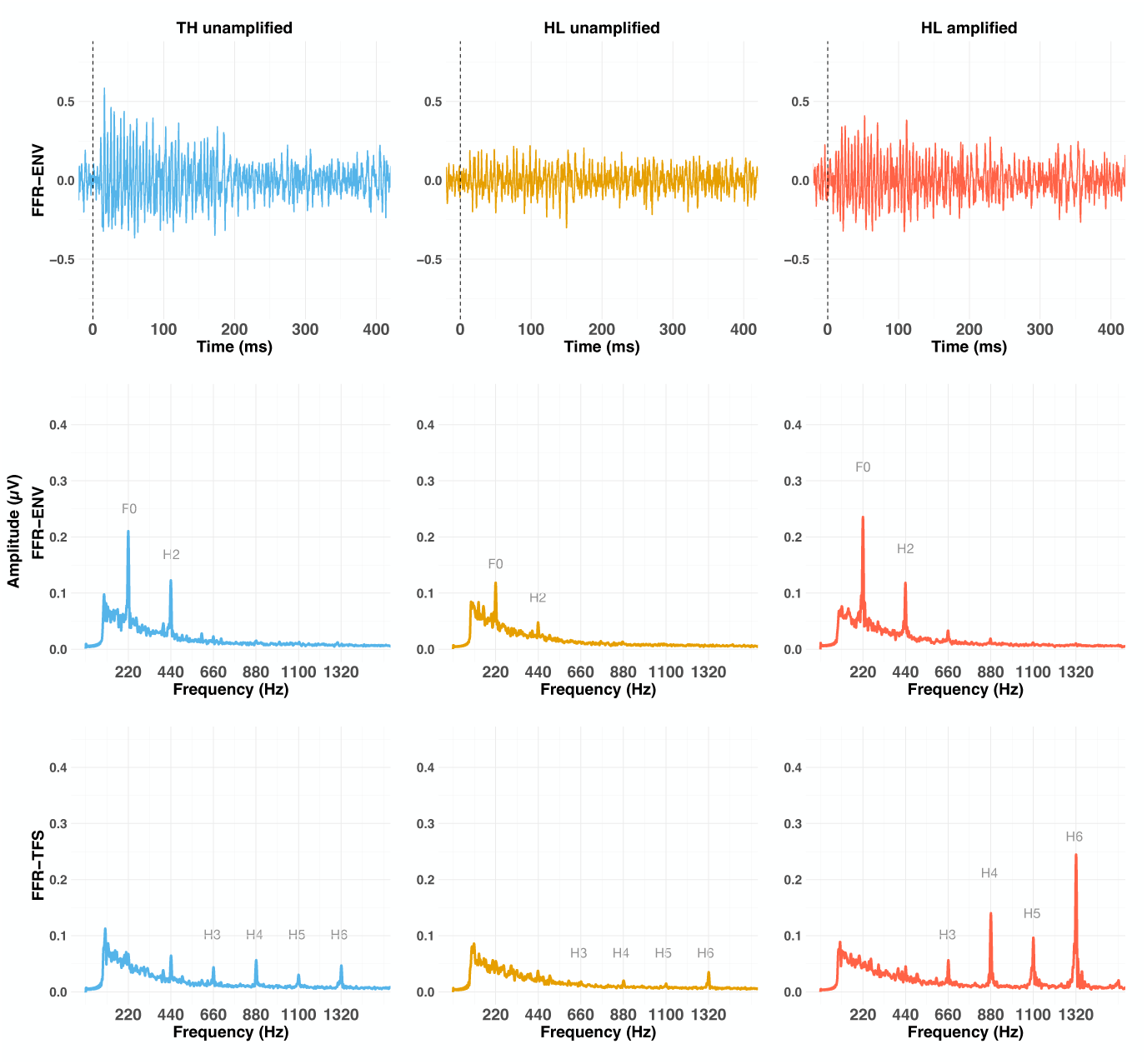
Grand average of the subcortical responses at Cz. The upper row represents the FFRENV in the time domain. The middle and lower rows represent the FFRENV and the FFRTFS evoked by the standard sound /b*ɑ*/, presented either unamplified to children with TH (70 dB SPL, left panel), unamplified to children with HL (70 dB SPL, middle column), or amplified to children with HL (with a frequency-specific gain tailored to each child’s hearing thresholds, right panel). The grand average spectra were computed by averaging the individual amplitude spectra.

Last, we compared the amplitude of the responses obtained when amplified sounds were presented to children with HL versus unamplified sounds presented to children with TH (HL_A_ vs. TH_U_; see Supplementary Table 3). The amplitude of the FFR_ENV_ did not significantly differ between children with HL_A_ and those with TH_U_ (*p* = 0.168). With respect to the FFR_TFS_, we observed a group × harmonic interaction (*p* < 0.001). FFR_TFS_ were significantly larger in children with HL_A_ than those with TH_U_ at both H4 and H6, but not at H3 and H5.

**Figure 4:**
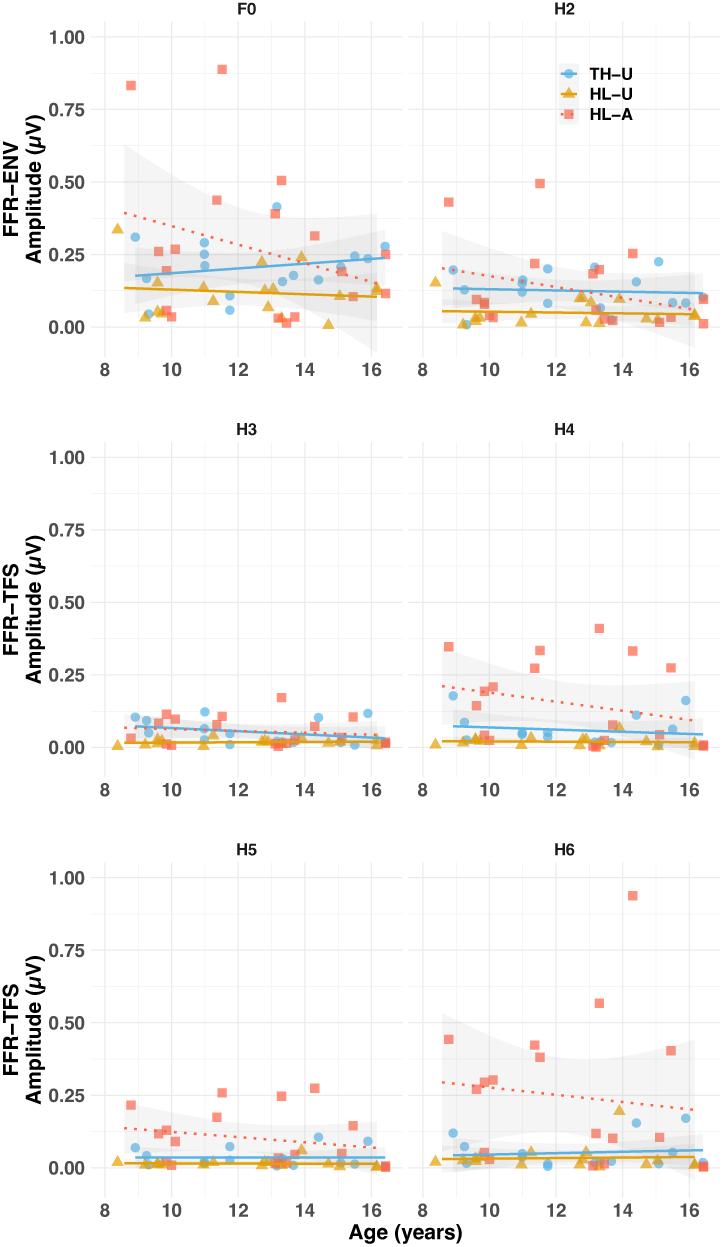
Scatterplots of the amplitude (µV) of the FFRENV (F0 and H2) and FFRTFS (H3 to H6) over age, for each group (THU, HLU and HLA). Regression lines are fit on the basis of each combination of group, FFR component and harmonic.

### 2. Cortical responses

**Figure 5:**
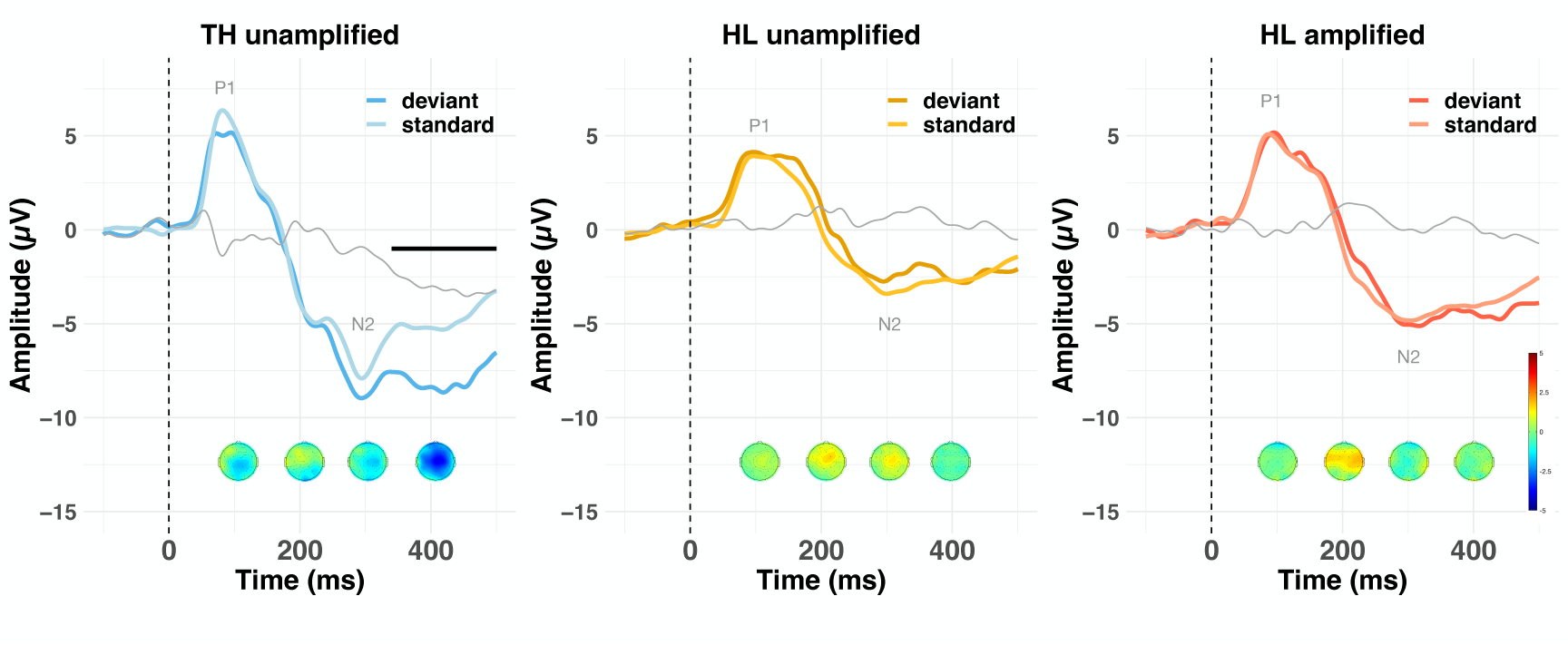
Grand average waveforms at Fz, evoked by the standard (lighter line) and deviant (darker line) presented either unamplified to both children with THU (70 dB SPL, left column) and children with HLU (70 dB SPL, middle column) or amplified to children with HLA (right column). The thin, gray line represents the differential wave between standards and deviants (MMN). Voltage maps show the mean MMN activity during the 100-500 ms post-stimulus time window. Negative values of the MMN are shown in blue, and positive values in red. The horizontal line represents the duration of a significant MMN in the period of the 100-500 ms post-stimulus-onset epoch evoked in children with TH. No significant MMNs were observed in either the unamplified or amplified conditions for the children with HL as a group.

Figure 5 shows the grand-average cortical waveforms evoked by standard and deviant stimuli at electrode Fz for both groups. In children with TH, the response to the standard sound was dominated by a large positivity occurring around 85 ms post-stimulus onset (the P1), followed by a large and prolonged negativity peaking around 300 ms (the N2). As we and others have reported previously, this pattern of results is typical of children with TH in this age range (Bishop et al., 2011; Calcus et al., 2019). For children with HL, the overall pattern of responses to the standard and deviant sounds was similar, albeit reduced in amplitude compared to children with TH. Note that there is no reliable evidence of an MMN in children with HL in either unamplified or amplified conditions, but strong evidence for one in children with TH (illustrated by a horizontal line covering the range from 330 to 500 ms in the left panel of Figure 5).

First, we assessed group differences in the cortical processing of unamplified speech (TH_U_ vs. HL_U;_ see Supplementary Table 4). As shown in Figure 6, P1 was smaller (by 2.05 µV) and later (by 2.79 ms) in children with HL_U_ than those with TH_U_ (both *p*s < 0.05). N2 was also smaller (by 3.79 µV; *p* < 0.001) but not later in children with HL_U_ than those with TH_U_. Additionally, children with HL_U_ had significantly smaller MMN responses (by 3.47 µV) than children with TH_U_ (*p* = 0.003).

Next, we sought to determine whether amplification benefited the cortical processing of speech sounds in children with HL (HL_U_ vs. HL_A;_ see Supplementary Table 5). For P1 amplitude, we found a significant amplification × BEPTA interaction (*p* = 0.028; see Supplementary Figure 1). Perhaps not surprisingly, the P1 amplitude in children with better BEPTA thresholds was not changed by the relatively small amplification they received, whereas the P1 amplitude for children with poorer BEPTA thresholds (and who received greater degrees of amplification) was increased so as to make P1 amplitude more-or-less independent of BEPTA. P1 latency was longer in the unamplified than the amplified condition (*p* = 0.027). It was also longer for individuals with higher BEPTA thresholds (*p* = 0.024) and for younger children (*p* = 0.005), irrespective of the condition for both. For N2 latency, there was a significant main effect of amplification (*p* = 0.033). Even though the amplification × BEPTA interaction was also significant (*p* = 0.030), post-hoc contrasts were not significant (*p* = 0.842). There was no significant effect of amplification on MMN amplitude or latency (both *p*s > 0.10). Note that irrespective of the condition, MMN amplitude decreased with age in children with HL (*p* = 0.040).

**Supplementary Figure 1:**
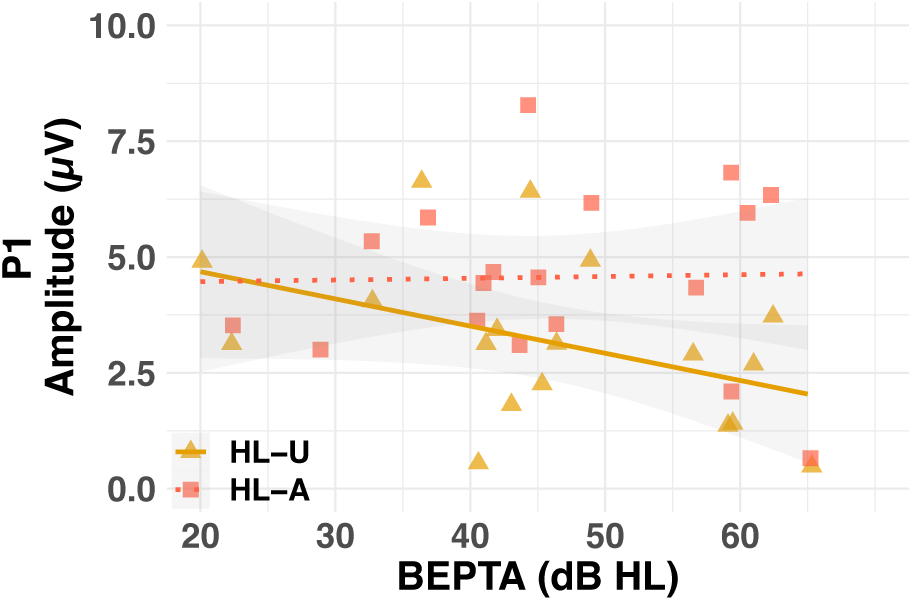
Amplitude (µV) of the P1 for the HL group as a function of BEPTA. Individual (shapes) and group (lines) data are shown for both unamplified (HLU, plain lines) and amplified (HLA, dotted lines) raw. Shaded lines represent the 95% confidence intervals.

Last, we evaluated the potential benefit of amplification for cortical processing of speech in children with HL to that of children with TH (HL_A_ vs. TH_U;_ see Supplementary Table 6). There was no significant effect of group on P1 amplitude or latency (both *p*s > 0.10). Yet there remained a significant group difference when looking at N2. Despite amplification, N2 amplitude was smaller (by 2.43 µV; *p* = 0.004), but not later, in the group with HL_A_ than the group with TH_U._ Similarly, children with HL_A_ showed significantly smaller MMN than the group with TH_U_ (*p* = 0.002). Note that, although the group × age interaction did not come out significant (*p* = 0.146), post-hoc comparisons indicate a significant decrease of MMN amplitude with age in children with HL_A_ (*p* = 0.038) but not in children with TH_U_ (*p* > 0.10).

**Figure 6:**
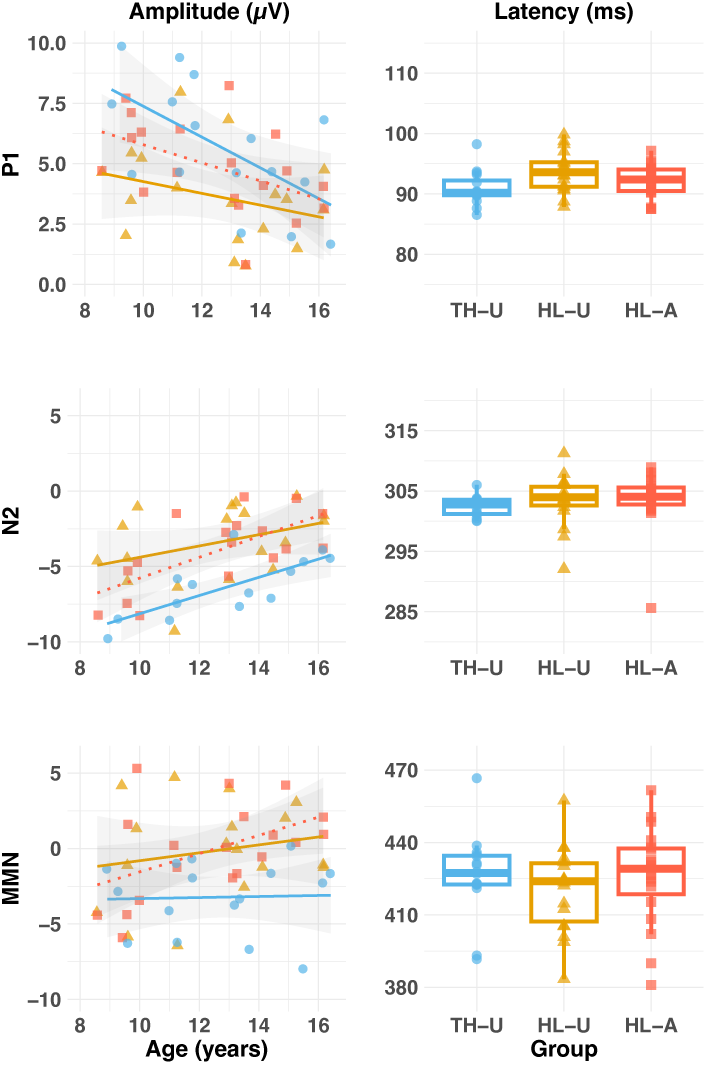
Amplitude (µV; left column) and latency (ms; right column) of the P1, N2 and MMN for each group and condition. Individual (shapes) and group (colour) data are shown for both unamplified (HLU, orange) and amplified (HLA, red) data. Shaded lines represent the 95% confidence intervals. Latencies (right column) are illustrated as group boxplots because there was no effect of age.

### 3. Behavioural data

#### 3.1. Speech discrimination in quiet

Figure 7 shows the behavioural performance of the two groups on the speech perception tasks. For speech discrimination of the speech sounds /bɑ/ vs /dɑ/ in quiet (Figure 7A), there was no significant effect of group, age, or their interaction on response thresholds (all *p*s > .10; see Supplementary Table 7A).

**Figure 7:**
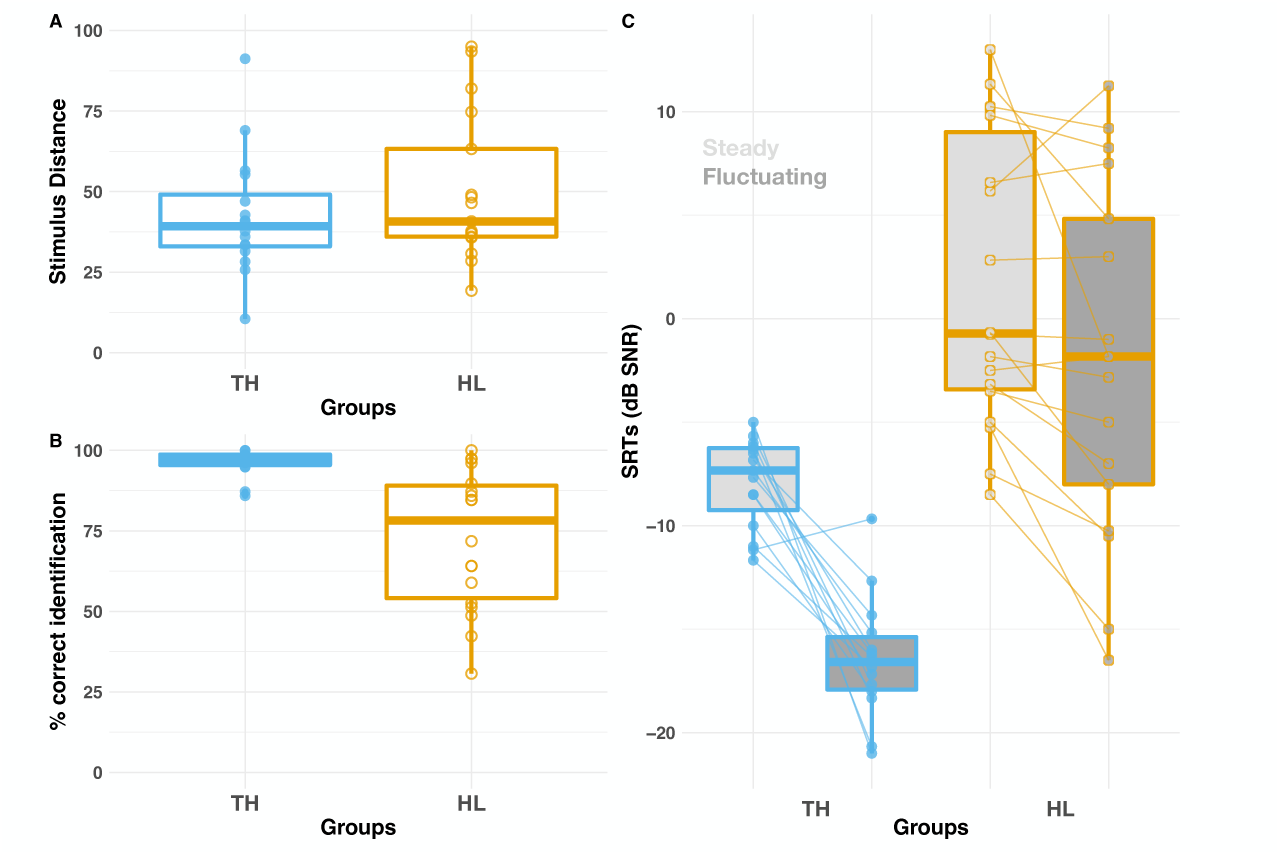
Boxplots of (A) stimulus distance (B) percentage correct consonant identification in quiet and (C) SRTs for consonant identification in steady (light grey) and fluctuating (dark grey) background noises. Children with TH are shown in blue, HL children are shown in orange. The boxes show the interquartile range (25^th^-75^th^ percentile) of the data with horizontal line indicating the median. The whiskers indicate values that fall within 1.5 times the interquartile range.

#### 3. 2. Speech identification in quiet and noise

Figure 7B presents the percentage correct consonant identification in quiet for the two groups (also see Supplementary Table 7B). Children with HL were significantly poorer than children with TH (*p* < 0.001). Note that for both groups, performance significantly improved with age (*p* = .003), although ceiling effects meant only a very small increase in percentage correct for the TH group.

Finally, Figure 7C presents the SRTs in steady and fluctuating background noise for the two groups (also see Supplementary Table 7C). The significant main effects of group and noise type (both *p*s < .001) need to be considered in light of the significant group × noise type interaction (*p* = .006). As expected, both groups experienced masking release when they were presented with fluctuating noise (both *p*s < 0.001). However, the magnitude of this effect was larger in children with TH (8.5 dB SNR) than in children with HL (3.9 dB SNR).

### 4. Relationship between the neural and behavioural responses

The relationship between electrophysiological activity and behavioural responses was investigated by means of a set of correlations. Holm-Bonferroni correction was applied to correct for multiple comparisons where appropriate.

Given the evidence that perception of speech in quiet is dominated by the encoding of the envelope of the signal, we investigated the relationship between the amplitude of the FFR_ENV_ at both F0 and H2 and behavioural performance of speech identification in quiet, separately for children with TH_U_ and children with HL_U_. None of the correlations were significant (all *p*s > 0.05).

Adequate perception of speech in noise requires precise spectro-temporal processing of speech, particularly in the dips of a fluctuating background noise. In fact, the amplitude of speech-evoked FFR in quiet relates to the perception of speech in noise (see introduction). Therefore, we investigated the relationship between amplitude of the speech-evoked FFR_TFS_ at H4 and H6 (in quiet) and behavioural speech perception in steady and fluctuating noise, as well as masking release, separately for children with TH and children with HL (in both unamplified and amplified conditions). None of the correlations were significant (all *p*s > 0.05).

The speech stimuli used for the electrophysiological recordings were the endpoints /bɑ/ and /dɑ/ of the continuum used for the speech discrimination in quiet behavioural task. As such, we hypothesised that a relationship might be observed between the amplitude of the MMN and the stimulus distance threshold measured in the speech discrimination in quiet behavioural task. However, there was no significant correlation between speech discrimination in quiet and MMN amplitude, as evaluated separately for both groups, and separately for the amplified and unamplified conditions in children with HL (all *p*s > 0.05).

Last, we investigated the relationship between subcortical and cortical responses, separately for both groups (and separately for HL_A_ and HL_U_). In children with TH, larger FFR_ENV_ at F0 were associated with smaller N2 responses (*p* = 0.004). There was no other significant correlation between FFR_ENV_ at F0 and P1, N2 or MMN (all *p*s > 0.05).

## Discussion

This study investigated the effect of childhood MM HL and the benefit of amplification on both subcortical and cortical processing of speech sounds. When sounds were presented at the same intensity across groups, both subcortical and cortical responses were smaller in children with HL than those with TH. When sounds were amplified, the subcortical responses increased in children with HL_A_ compared to HL_U_. In fact, they could even be larger in children with HL_A_ than TH_U_ (although only significantly so for the FFR_TFS_). At the cortical level, an amplification benefit was also observed on P1 amplitude but not on the amplitude of later responses (N2 or MMN). In fact, at the group level, children with HL did not show a significant MMN (unlike children with TH), whether the sounds were amplified or not. When looking at individual responses, the amplitude of the MMN tended to decrease with age in children with HL but not in those with TH. Behaviourally, children with HL had poorer speech intelligibility than those with TH both in quiet and in noise, and they showed less benefit from background noise fluctuations. Yet, the relationship between the neurophysiological responses and the behavioural performance remains unclear.

### Subcortical processing

When presented with unamplified sounds, children with MM HL showed significantly smaller FFR_ENV_ and FFR_TFS_ than those with TH. This finding replicates and extends previous results from older adults with age-related HL (Plyler & Ananthanarayan 2001; Ananthakrishnan et al. 2016) – but contrasts with some studies that did not find group differences on unamplified FFR measures (Anderson et al. 2013; Presacco et al. 2019). The etiology of the HL might thus modulate its effect on subcortical processing of unamplified sounds. When presented with amplified sounds, children with MM HL showed a clear benefit from the frequency-specific gain applied to the sounds. In fact, the amplified FFR_ENV_ of children with MM HL did not significantly differ from that of children with TH_U_. Furthermore, the amplified FFR_TFS_ was even larger than that of children with TH_U_. Our findings thus fit with existing results from the adult and child literature to indicate a clear benefit of amplification at the subcortical level (Anderson et al., 2013; BinKhamis et al., 2019; Easwar et al., 2015, 2023, 2024; Jenkins et al., 1993).

We were initially surprised by the spectacular increase in amplitude of the FFR_TFS_ in the higher harmonics, especially H6, following amplification. However, it is crucial to keep in mind that the components reflected in this spectrum are not directly present on the scalp-recorded EEG, but arise through a subtraction of the waves of the two stimulating polarities. Consider Supplementary Figure 2, which depicts the spectra of the positive and negative polarity stimuli separately. For children with HL_A_, the spectra do indeed show a large component at H6 (the largest component in the FFR_TFS_), but one which is roughly the size of the one at F0. This is not such a different pattern from that shown by children with TH_U_, although the F0 is somewhat larger than H6 in this latter case. The large differences between these two groups in the FFR_TFS_ spectra arise not only from the component spectral amplitudes, but also because the *phases* of the original components change over frequency. F0 and H2 have very similar phases in the negative and positive polarities (thus leading to their enhancement in the FFR_ENV_ spectra and cancellation in the FFR_TFS_ spectra), whereas, H3-H6 have phases that are close to 180° out of phase with one another (thus leading to their cancellation in the FFR_ENV_ spectra and enhancement in the FFR_TFS_ spectra). In that respect, the size of the higher frequency components is roughly doubled in the FFR_TFS_ spectra, as compared to their actual measured sizes in the individual polarity waves.

We can also exclude the possibility of electrical artefacts from the stimuli by noting that the latency of the FFR_TFS_ and FFR_ENV_ responses for all groups was very similar, even though any artefacts would be cancelled in the FFR_ENV_ responses. Note too that high amplitudes of high harmonics have been seen before in FFR_TFS_ spectra. Krishan (2002), in particular, for a similar /ɑ/ vowel to the one used here (but based on a male voice with F0 ≈ 125 Hz), found, at the highest level tested, that the largest spectral components was at H7 (very close to the first formant frequency).

We believe these high amplitudes of H4-H6 (at 880 - 1320 Hz) stem from applying the individualized, frequency-specific NAL gains calculated from audiograms with sloping hearing losses (leading to increasing gains for higher frequency harmonics), but also because the vowel has a broad spectral peak in that region (see Fig 2) due to the F1 and F2 frequencies of 800 and 1340 Hz. To our knowledge, the only other study^47^ that used the NAL formula analysed FFR_TFS_ responses only up to ∼700 Hz. In this frequency range, there was no significant difference in the FFR_TFS_ of older adults with HL (in quiet, with or without amplification) and age-matched adults with TH. In our study, FFR_TFS_ was only above the noise floor at frequencies at H3 (660 Hz) and above. In fact, our results show significantly larger FFR_TFS_ in children with HL_A_ compared to both children with HL_U_ and children with TH_U_ at H4 and H6, which are close to the first and second formant peaks of our stimulus. In adults with TH, FFR_TFS_ are most clearly detected at harmonics that are close to formant peaks, up to 1500 Hz^24^. In short, especially large FFR_TFS_ at H4 - H6 might therefore reflect the interaction between frequency-specific NAL amplification and the formants of our speech stimuli. Yet a *larger* FFR_TFS_ in children with HL_A_ than those with TH_U_ might indicate hyperactivity at the subcortical level following childhood HL.

Hyperactivity has been widely reported as a consequence of SNHL (for a review, see Herrmann & Butler, 2021), and is thought to compensate for reduced inputs from damaged peripheral structures to central brain regions, to maintain sensation (Chambers et al., 2016). Such compensation fits within the theoretical framework of homeostatic plasticity, which states that neurons regulate their physiological properties to compensate for persistent changes in their activity level (Turrigiano & Nelson, 2004). Homeostatic mechanisms are thought to operate from synaptic to network levels of sensory systems. That FFR amplitude was larger in children with HL than those with TH, but only when sounds were amplified suggests that compensatory activity is intensity-dependent at the brainstem level. In mice, changes in auditory cortex responsiveness following conductive HL were only observed in sounds above a certain threshold, or in certain frequency ranges (Teichert et al., 2017). Future studies are warranted to investigate the effect of audibility on homeostatic plasticity of subcortical processing.

### Cortical processing

Despite the medium to large benefit of amplification on subcortical responses, there was only a small benefit of amplification on cortical responses (P1), or no benefit at all (N2, MMN). Individualized frequency-specific gain increased P1 amplitude (which reflects improved sound detectability, Eggermont & Ponton 2002), but only for children with poorer BEPTA. Irrespective of BEPTA, P1 was earlier in HL_A_ than HL_U_. In fact, the P1 amplitude and latency of HL_A_ did not significantly differ from that of children with TH_U_. This is in line with studies that show age-appropriate P1 when sounds are presented at similar sensation level to children with HL and those with TH (Koravand et al. 2013; Martinez et al. 2013; Engström et al. 2021).

Despite preserved P1, our results indicate that childhood HL leads to smaller (but not later) N2 even when sounds are amplified, in line with Koravand et al., 2013, but contrary to Martinez et al., 2013 or Engström et al., 2021. Furthermore, we did not observe a significant MMN at the group level on children with HL, even when sounds were amplified. At the individual level, the MMN amplitude of children with HL_A_ remained smaller than those with TH_U_. Note however that there was no group difference in MMN latency. Our finding of absent/smaller MMNs in 8 to 16 year-olds with HL contrast with those of age-appropriate MMNs in 5-12 year-old children (Koravand et al., 2013; Engström et al., 2021). Part of the explanation for this discrepancy might lie in the developmental effects of childhood HL on the central auditory pathway (see below for a discussion of the developmental effects observed in this study).

Why does the P1 amplitude benefit from amplification, but not the N2 or MMN? One possible explanation might lie in the different generators of the P1, N2 and MMN. The P1 generator has been reliably located within the secondary auditory cortex (Godey et al., 2001; Liegeois-Chauvel et al., 2004), whereas MMN generators likely include an extended thalamocortical circuitry projecting to primary auditory cortex (Lakatos et al., 2020). Little is known about the generators of the N2, but early intracranial recordings also seem to point towards a subcortical (thalamic) source (Velasco et al., 1989). Although anatomically close, the primary and secondary auditory cortices receive input from distinct portions of the medial geniculate body. Whereas the lemniscal pathway connects the ventral medial geniculate body to the primary auditory cortex, the nonlemniscal pathway connects the medial/dorsal geniculate body to both primary and secondary auditory cortices (Webster, 1992). Given that our results indicate smaller N2 and MMN even when children with HL were presented with amplified sounds, these could indicate a specific alteration of the nonlemniscal pathway connecting the medial portion of the geniculate body to the primary auditory cortex. Our cortical findings thus contrast with the subcortical evidence of hyperactivity following childhood HL.

### The effect of HL on the development of the central auditory pathway

In fact, our cortical findings also contrast with findings of hyperactivity in the auditory cortex as a consequence of age-related HL. Indeed, studies have reported some degree of enhancement of the cortical responses to sounds in older adults with age-related HL compared to age-matched individuals with TH (Alain et al., 2014; Tremblay et al., 2003; Millman et al., 2017). To our knowledge, none of the pediatric studies investigating cortical processing of (amplified or unamplified) sounds in children with HL found such indications of increased cortical responsiveness (Calcus et al., 2019; Engström et al., 2021; Ji et al., 2023; Martinez et al., 2013; Koravand et al., 2013; Rance et al., 2002; Glista et al., 2012; Uhler et al., 2023). In mice, the same population of neurons showed distinct mechanisms of homeostatic adaptation if visual deprivation occurred in juvenile or adult animals (Wen & Turrigiano, 2021). Such an age-dependent effect of homeostatic plasticity might also be observed in the central auditory pathway following auditory deprivation.

Noteworthily, these age-dependent effects might already be operating between 8 and 16 years of age. Indeed, we observed a significant decrease in the benefit of amplification with age on FFR_ENV_ amplitude. Additionally, the amplitude of the MMN decreased with age in children with HL_A_ (but not in those with TH_U_) – although note that the group × age interaction was not significant. Both results seem to indicate that adolescence could be a period of heightened plasticity for functional changes following auditory deprivation. Future studies are warranted to investigate this claim in a larger sample size, ideally taking puberty into account, as it could act as a biological trigger to those alterations (Gracia-Tabuenca et al., 2021; Laube et al., 2020; Vijayakumar et al., 2021).

### Multiple gain mechanisms within the auditory system

To our knowledge, only three studies so far have combined subcortical and cortical measures from the same individuals to investigate the effects of HL on auditory processing. Speech-evoked subcortical and cortical responses were recorded successively in older adults, with and without hearing aids. When sounds were presented at 65 dB SPL, hearing aid amplification lead to significantly larger subcortical, but not cortical responses (Jenkins et al, 2017). In gerbils with developmental conductive HL, age-appropriate subcortical, but not cortical responses were recorded to amplitude modulated sounds presented at similar sensation level as in animals with TH (Yao & Sanes, 2018). More recently, Hutchison et al. (2023) recorded subcortical and cortical neurophysiological responses in young adults with TH before, throughout and after fitting them with earplugs. The results indicate an increase in the subcortical response, but a reduction of cortical activity ipsilateral to the deprived ear. Together with our observations, the literature suggests that multiple gain mechanisms might underlie homeostatic plasticity throughout the auditory pathway (Munro et al., 2014). Neurons from each level of the auditory pathway are thought to display idiosyncratic changes following HL (for a review, see Sanes, 2013).

### Relationship between speech processing & perception

An exploratory objective of our study was to investigate how changes in neurophysiological responses following HL might relate to speech intelligibility in quiet and in noise. To do so, we chose stimuli that were as similar as possible for both neurophysiological and behavioural tasks. However, two methodological choices are worth mentioning here: Unlike the neurophysiological task, stimuli were only presented unamplified in the behavioural tasks. Additionally, syllables were only presented in quiet in the neurophysiological tasks; yet speech perception was investigated in quiet and in noise.

Behaviourally, children/adolescents with HL have higher (i.e. poorer) thresholds in noise than children/adolescents with TH (Lewis et al. 2015; Goldsworthy & Markle 2019). Their thresholds improved in the presence of fluctuating compared to steady background noise, but to a lesser extent than the group with TH. Similarly to adults with HL (Lorenzi et al. 2006; Hopkins et al. 2008; Strelcyk & Dau 2009), children with MM HL seem less able to listen in the “valleys” of a fluctuating background noise to improve their speech perception. However, we did not find any significant relationship between measures of speech processing and perception. This contrasts with numerous studies that report small to medium correlations between subcortical responses and speech intelligibility, especially in noise (Anderson et al., 2010; Bidelman & Momtaz, 2021; Hornickel et al., 2009; Mai et al., 2018; Mepani et al., 2021; van Hirtum et al., 2023). Some correlations have also been observed between cortical responses and speech discrimination (Ching et al., 2023; Näätänen, 2001), although the nature of this relationship remains unclear (Jost et al., 2015; Sharma et al., 2006; Shafer et al., 2005). Again, our sample size might be too small to adequately investigate such possible correlations. Future studies including larger sample sizes are warranted to clarify the relationship between speech processing and perception in the context of congenital HL. These studies would benefit from a direct investigation of the effect of childhood HL on subcortical processing of speech in noise and its relationship with performance.

## Conclusions

When presented with unamplified sounds, children with HL show impaired subcortical and cortical responses to speech syllables. Amplification was beneficial for subcortical and early (P1) but not late (N2, MMN) cortical responses. Our results support the existence of multiple gain mechanisms that could underlie homeostatic plasticity throughout the auditory pathway. Responses elicited at different levels of the auditory pathway appear to display idiosyncratic changes following HL, and may be age- and level-dependent. Even though speech perception was poorer in the group with HL than TH (in quiet and in noise), there was no clear relationship between the neurophysiological and behavioural measures. Even mild to moderate congenital HL induces functional alterations to the central auditory pathway, each level showing a specific alteration of its typical response. Future studies are needed to clarify the relationship between these functional changes and the behavioural performance.

**Supplementary Figure 2:**
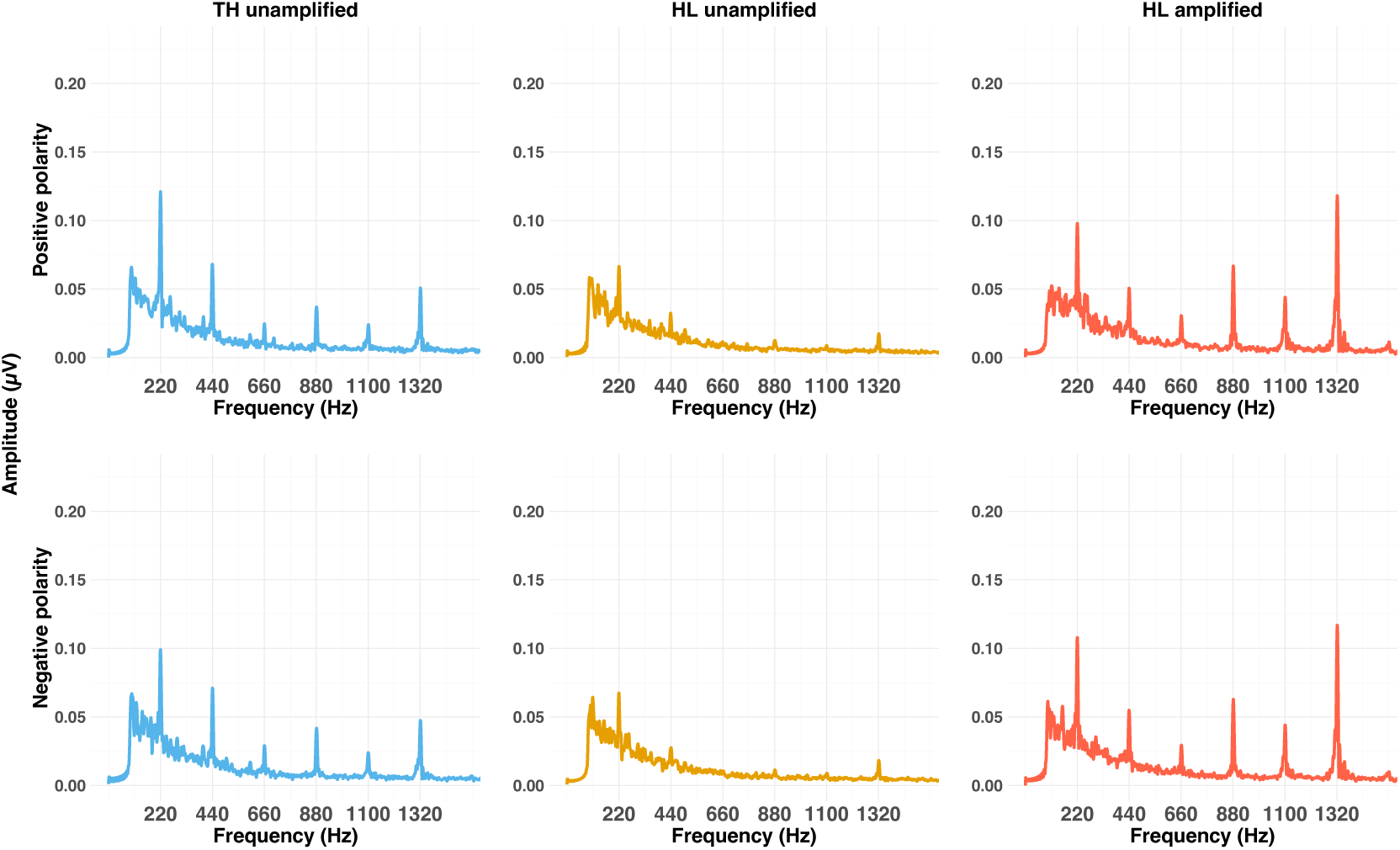
Grand average of the subcortical responses at Cz, as elicited by stimulus in each polarity. The upper row represents the spectra evoked by the standard syllable /bɑ/, presented in positive polarity; the lower row represents the spectra evoked by the same stimulus presented in negative polarity. Grand average spectra were computed by averaging the individual amplitude spectra.

## Acknowledgments

We are very grateful for the time and efforts of the children who participated in this study and to their parents. We would like to thank Dr Lorna Halliday for her input throughout this project, and for her comments on previous versions of this manuscript. We would also like to thank Gaston Hilkhuysen for his help with aspects of the methods. We thank two anonymous reviewers for their constructive feedback on an earlier version of this manuscript. This work was supported by the FP7 people programme (Marie Curie Actions), grant agreement no FP7-607139 (improving Children’s Auditory REhabilitation, iCARE). Axelle Calcus was supported by a MSCA Individual Fellowship (798093, EAR-DNA) and a FNRS MIS (F.4508.22).

## Author contributions

A.C. and S.R. conceived the idea, A.C. collected data, A.C. and S.R. analysed the data, A.C. and S.R. interpreted the data, A.C. and S.R. wrote the manuscript.

## Data availability

Unidentifiable data, stimuli and statistical analyses scripts are publicly available on: https://github.com/acalcus/MMHL_amplification

## Competing interests

The authors declare no competing interests

## Notes

https://github.com/acalcus/MMHL_amplification

### Competing Interest Statement

The authors have declared no competing interest.

### Summary of Updates

The introduction has been re-worked to better distinguish the exploratory analyses from the confirmatory hypotheses. Statistical analyses have been slightly modified to better account for the predictors we used. Further analyses were performed to better understand the large FFR-TFS responses were recorded. The discussion has been updated accordingly to the changes made to the responses.

