## Supplemental table1 for "Effects of childhood hearing loss on the subcortical and cortical representation of speech"

**Supplementary Table 1:** Final model of the effects of hearing loss on the subcortical responses: between-participants ($NH_U$ vs. $HL_U$) comparisons.

| Dependant variable | Regressor | *Coefficient* | *Std error* | *df* | *t* | *p* | $\eta^2$ |
|---|---|---|---|---|---|---|---|
| $FFR_{ENV}$ | **Group** | **0.082** | **0.022** | **31** | **3.61** | **0.001** | **0.20** |
| | **Harmonic** | **-0.076** | **0.012** | **32** | **-6.13** | **< 0.001** | **0.17** |
| $FFR_{TFS}$ | **Group** | **0.030** | **0.012** | **29.0** | **2.57** | **0.015** | **0.13** |
| | **Harmonic** (H3 vs H6) | **0.010** | **0.005** | **89.7** | **1.93** | **0.056** | **-** |

Note: The best fitting model for $FFR_{ENV}$ and $FFR_{TFS}$ were all an acceptable fit ($FFR_{ENV}$ [AIC = -140.66, $R^2c$ = 0.706], $FFR_{TFS}$ [AIC = -494.49, $R^2c$ = 0.767]). Significant results are shown in boldface.

**Supplementary Table 2:** Final model of the effects of amplification on the subcortical responses: within-participants comparisons ($HL_U$ vs. $HL_A$).

| Dependant variable | Regressor | *Coefficient* | *Std error* | *df* | *t* | *p* | $\eta^2$ |
|---|---|---|---|---|---|---|---|
| $FFR_{ENV}$ | **Amplification** | **-0.119** | **0.042** | **17.2** | **-2.83** | **0.011** | **0.11** |
| | **Harmonic** | **-0.106** | **0.018** | **50.0** | **-5.81** | **< 0.001** | **0.09** |
| $FFR_{TFS}$ | Amplification | -0.039 | 0.035 | 36.7 | -1.11 | 0.27 | 0.19 |
| | **Harmonic** (H3 vs H4) | **0.097** | **0.023** | **110.9** | **4.24** | **< 0.001** | **-** |
| | **Harmonic** H3 vs H5 | **0.047** | **0.023** | **110.9** | **2.06** | **0.041** | **-** |
| | **Harmonic** H3 vs H6 | **0.192** | **0.023** | **110.9** | **8.40** | **< 0.001** | **-** |
| | **Amplification × Harmonic** H4: $HL_U$ vs $HL_A$ | **0.134** | **0.035** | **36.4** | **3.79** | **< 0.001** | **-** |
| | **Amplification × Harmonic** H5: $HL_U$ vs $HL_A$ | **0.089** | **0.035** | **36.4** | **2.53** | **0.016** | **-** |
| | **Amplification × Harmonic** H6: $HL_U$ vs $HL_A$ | **0.215** | **0.035** | **36.4** | **6.07** | **< 0.001** | |

Note: The best fitting model for $FFR_{ENV}$ and $FFR_{TFS}$ were all an acceptable fit ($FFR_{ENV}$ [AIC = -85.52, $R^2c$ = 0.812], $FFR_{TFS}$ [AIC = -231.05, $R^2c$ = 0.573]).

**Supplementary Table 3:** Final model of the effects of hearing loss on the subcortical responses: between-participants ($NH_U$ vs. $HL_A$) comparisons.

| Dependant variable | Regressor | *Coefficient* | *Std error* | *df* | *t* | *p* | $\eta^2$ |
|---|---|---|---|---|---|---|---|
| $FFR_{ENV}$ | **Harmonic** | **-0.113** | **0.020** | **33** | **-5.67** | **< 0.001** | **0.11** |
| $FFR_{TFS}$ | Group | -0.003 | 0.040 | 60.4 | -0.08 | 0.932 | 0.01 |
| | **Harmonic** H3 vs H4 | **0.097** | **0.024** | **93.3** | **3.93** | **<0.001** | **-** |
| | **Harmonic** H3 vs H6 | **0.192** | **0.024** | **93.3** | **7.78** | **<0.001** | **-** |
| | **Group × Harmonic** H4: $HL_A$ vs $TH_U$ | **0.091** | **0.041** | **61.7** | **2.23** | **0.029** | **-** |
| | **Group × Harmonic** H6: $HL_A$ vs $TH_U$ | **0.192** | **0.041** | **61.7** | **4.70** | **<0.001** | **-** |

Note: The best fitting model for $FFR_{ENV}$ and $FFR_{TFS}$ were all an acceptable fit ($FFR_{ENV}$ [AIC = -64.21, $R^2c$ = 0.766], $FFR_{TFS}$ [AIC = -191.79, $R^2c$ = 0.706]).

**Supplementary Table 4:** Final model of the effects of hearing loss on the cortical responses: between-participants comparisons ($HL_{U\ vs.}$ $NH_U$). Significant predictors are shown in boldface.

| Dependant variable | Regressor | *Coefficient* | *Std error* | *df* | *t* | *p* | $\eta^2$ |
|---|---|---|---|---|---|---|---|
| P1 amplitude | **Group** | **2.088** | **0.701** | **31** | **2.97** | **0.005** | **0.18** |
| | **Age** | **-0.427** | **0.148** | **31** | **-2.88** | **0.007** | **0.17** |
| P1 latency | **Group** | **-2.798** | **1.065** | **32** | **-2.62** | **0.013** | **0.17** |
| N2 amplitude | **Group** | **-3.835** | **0.814** | **31** | **-4.71** | **< 0.001** | **0.34** |
| | **Age** | **0.615** | **0.172** | **31** | **3.57** | **0.001** | **0.19** |
| N2 latency | - | - | - | - | - | - | - |
| MMN amplitude | **Group** | **-3.480** | **1.085** | **32** | **-3.20** | **0.003** | **0.24** |
| MMN latency | - | - | - | - | - | - | - |

Note: The best fitting model for P1, N2 and MMN amplitude and latency were all an acceptable fit (P1 amplitude [AIC = 149.88, $R^2$ = 0.353], P1 latency [AIC = 177.38, $R^2$ = 0.177], N2 amplitude [AIC = 159.99, $R^2$ = 0.526], N2 latency [AIC = 180.61, $R^2$ = 0.025], MMN amplitude [AIC = 178.63, $R^2$ = 0.243], MMN latency [AIC = 294.49, $R^2$ = 0.031]).

**Supplementary Table 5:** Final model of the effects of hearing loss on the cortical responses: within-participants comparisons ($HL_U$ vs. $HL_A$). Significant predictors are shown in boldface.

| Dependant variable | Regressor | *Coefficient* | *Std error* | *df* | *t* | *p* | $\eta^2$ |
|---|---|---|---|---|---|---|---|
| P1 amplitude | Amplification | 1.616 | 1.238 | 16.4 | 1.30 | 0.210 | 0.08 |
| | BEPTA | 0.000 | 0.031 | 21.1 | 0.00 | 0.998 | 0.04 |
| | **Age** | **-0.331** | **0.156** | **16.0** | **-2.12** | **0.05** | **0.17** |
| | **Amplification × BEPTA** | **-0.063** | **0.026** | **16.3** | **-2.41** | **0.028** | **0.05** |
| P1 latency | **Amplification** | **1.639** | **0.678** | **17.5** | **2.41** | **0.026** | **0.06** |
| | **Age** | **-0.605** | **0.188** | **16.1** | **-3.21** | **0.005** | **0.20** |
| | **BEPTA** | **0.086** | **0.035** | **16.3** | **2.48** | **0.024** | **0.13** |
| N2 amplitude | **Age** | **0.682** | **0.211** | **16.9** | **3.24** | **0.004** | **0.28** |
| N2 latency | **Amplification** | **-6.421** | **2.768** | **16.6** | **-2.32** | **0.033** | **0.00** |
| | BEPTA | -0.087 | 0.080 | 21.5 | -1.091 | 0.287 | 0.00 |
| | **Amplification × BEPTA** | **0.138** | **0.058** | **16.5** | **2.37** | **0.030** | **0.04** |
| MMN amplitude | **Age** | **0.506** | **0.228** | **17.0** | **2.22** | **0.040** | **0.13** |
| MMN latency | - | | | - | - | - | - |

Note: The best fitting model for P1, N2 and MMN amplitude and latency were all an acceptable fit (P1 amplitude [AIC = 157.50, $R^2c$ = 0.768], P1 latency [AIC = 181.40, $R^2c$ = 0.593], N2 amplitude [AIC = 182.70, $R^2c$ = 0.499], N2 latency [AIC = 213.64, $R^2c$ = 0.768], MMN amplitude [AIC = 196.83, $R^2c$ = 0.200], MMN latency [AIC = 323.05, $R^2c$ = 0.109]).

**Supplementary Table 6:** Final model of the effects of hearing loss on the cortical responses: between- participants comparisons ($HL_A$ vs. $NH_U$). Significant predictors are shown in boldface.

| Dependant variable | Regressor | *Coefficient* | *Std error* | *df* | *t* | *p* | $\eta^2$ |
|---|---|---|---|---|---|---|---|
| P1 amplitude | **Age** | **-0.493** | **0.135** | **32** | **-3.65** | **< 0.001** | **0.28** |
| P1 latency | - | - | - | - | - | - | - |
| N2 amplitude | **Group** | **-2.622** | **0.846** | **32** | **-3.10** | **0.004** | **0.14** |
| | **Age** | **0.918** | **0.178** | **32** | **5.15** | **< 0.001** | **0.39** |
| N2 latency | - | - | - | - | - | - | - |
| MMN amplitude | **Group** | **-3.152** | **0.951** | **33** | **-3.31** | **0.002** | **0.26** |
| MMN latency | - | - | - | - | - | - | - |

Note: The best fitting model for P1, N2 and MMN amplitude and latency were all an acceptable fit (P1 amplitude [AIC = 148.81, $R^2$ = 0.334], P1 latency [AIC = 174.26, $R^2$ = 0.047], N2 amplitude [AIC = 168.11, $R^2$ = 0.521], N2 latency [AIC = 195.53, $R^2$ = 0.025], MMN amplitude [AIC = 175.45, $R^2$ = 0.250], MMN latency [AIC = 308.92, $R^2$ = 0.000]).

**Supplementary Table 7:** Group comparisons (NH vs. HL) on the behavioural data. Significant predictors are shown in boldface.

| Dependant variable | Regressor | *Statistic (F or z)* | *df* | *p* | $\eta^2$ |
|---|---|---|---|---|---|
| **A.** Stimulus distance (in quiet) | - | - | - | - | - |
| | - | - | - | - | - |
| **B.** % correct (in quiet) | **Group** | **-5.38** | | **< 0.001** | |
| | **Age** | **2.92** | | **0.003** | |
| **C.** SRTs (in noise) | **Group** | **38.26** | **(1, 30.9)** | **< 0.001** | **0.44** |
| | **Noise type** | **50.85** | **(1, 29.4)** | **< 0.001** | **0.11** |
| | **Group × Noise type** | **8.72** | **(1, 29.4)** | **0.006** | **0.02** |

Note: The best fitting model for the behavioural data were all an acceptable fit (**A.** stimulus distance [AIC = 302.613, $R^2$ = 0.042], **B.** % correct in quiet [AIC = 1700.9], **C.** SRTs in noise [AIC = 389.545, $R^2c$ = 0.868])
